# pHusion: A robust and versatile toolset for automated detection and analysis of exocytosis

**DOI:** 10.1101/2023.07.25.550499

**Authors:** Ellen C. O’Shaughnessy, Mable Lam, Samantha E. Ryken, Theresa Wiesner, Kimberly Lukasik, Bradley J Zuchero, Christophe Leterrier, David Adalsteinsson, Stephanie L. Gupton

## Abstract

Exocytosis is a fundamental process used by eukaryotic cells to regulate the composition of the plasma membrane and facilitate cell-cell communication. To investigate the role exocytosis plays in neuronal morphogenesis, previously we developed computational tools with a graphical user interface (GUI) to enable the automatic detection and analysis of exocytic events (ADAE GUI) from fluorescence timelapse images. Though these tools have proven useful, we found that the code was brittle and not easily adapted to different experimental conditions. Here, we have developed and validated a robust and versatile toolkit, named pHusion, for the analysis of exocytosis written in ImageTank, a graphical programming language that combines image visualization and numerical methods. We tested this method using a variety of imaging modalities and pH-sensitive fluorophores, diverse cell types, and various exocytic markers to generate a flexible and intuitive package. Using pHusion, we show that VAMP3-mediated exocytosis occurs 30-times more frequently in melanoma cells compared with primary oligodendrocytes, that VAMP2-mediated fusion events in mature rat hippocampal neurons are longer lasting than those in immature murine cortical neurons, and that exocytic events are clustered in space yet random in time in developing cortical neurons.

**Summary Statement:** Exocytosis is an essential process by which cells change shape, alter membrane composition, and communicate with other cells. Though all eukaryotic cells carry out exocytosis, the regulation of vesicle fusion, the cargo of vesicles, and the role exocytosis plays in cell fate differ greatly across cell types. Here, we developed a flexible and robust set of tools to enable automatic identification and analysis of exocytic events across a wide range of cell types, vesicle types, and imaging conditions.

## Introduction

Exocytosis is an essential biological process during which secretory vesicles fuse with the plasma membrane (PM) thereby altering the composition of the PM and releasing lumenal cargo into the extracellular space. Though common to eukaryotic cells, exocytosis serves myriad functions from altering membrane fluidity and changing cell shape to facilitating synaptic transmission, hormone release, and remodeling the extracellular matrix (Burgoyne and Morgan, 2003; Lin and Scheller, 2000; Mostov et al., 2000; Winkle et al., 2014). The soluble N-ethylmaleimide-sensitive factor attachment proteins receptors (SNARE) complex is the minimal protein machinery required for exocytosis (Block et al., 1988; Jahn and Fasshauer, 2012; Südhof and Rothman, 2009; Weber et al., 1998). To enable fusion of membranes, SNARE proteins attached to different membranes form a tightly packed bundle of four α-helical coiled-coils (CCs)(Gao et al., 2012; Söllner et al., 1993; Sutton et al., 1998). Vesicle SNAREs (vSNAREs), such as the vesicle associated membrane protein (VAMP) family, contribute one α-helix and plasma membrane-targeted SNAREs (tSNARES), such as the syntaxin1 and SNAP25 families of proteins, contribute the other three α-helices (Söllner et al., 1993). Which SNAREs are bundled in a SNARE complex is a tightly regulated process that depends heavily on the cell type, developmental stage, and exocytic mode (Urbina and Gupton, 2020). In neurons, two distinct modes of exocytosis have been elucidated: full vesicle fusion (FVF) in which membrane material is delivered to the plasma membrane and kiss-and-run (KNR) in which vesicular cargo is released into the extracellular environment with no membrane incorporation (Bowser and Khakh, 2007; Harata et al., 2001; Heuser and Reese, 1973; Urbina et al., 2021). Although this core machinery involved in vesicle fusion has been elucidated over the past 40 years, understanding how these diverse processes are regulated remains an active area of investigation.

Recent advances in microscopy and better fluorescent tools to visualize exocytosis have given rise to large, image-based, data sets poised to address previously inaccessible questions. Despite the availability of high quality imaging data, image analysis frequently presents a bottleneck to discovery. Manual image analysis is problematic as it is error prone, subject to bias, and time consuming. To overcome these limitations in our work studying neuronal morphogenesis, we previously developed ADAE GUI, a semi-automated computer-vision based toolkit using MATLAB (The MathWorks Inc., 2021) and R (R: The R Project for Statistical Computing, 2021) to identify exocytic events, characterize their spatio-temporal dynamics and classify distinct modes of exocytosis (Urbina and Gupton, 2021; Urbina et al., 2018; Urbina et al., 2021).

Though these computational tools were instrumental in several studies (Lam et al., 2022; Urbina et al., 2018; Urbina et al., 2021; Ye et al., 2022), we have found that different modes of microscopy, such as total internal reflection fluorescence (TIRF) vs. widefield epifluorescence microscopy, different camera sensor technologies, tagging different vSNARE proteins with the commonly used fluorophore, pHluorin, or use of alternate pH-sensitive fluorescent proteins greatly impacted the functionality of this code. Unfortunately the intermediate steps in ADAE GUI are opaque, both in terms of the code and the inability to visualize image transformations, making it challenging to identify and address points of failure. In addition, collaborators studying disparate cell types and/or distinct stages of neuronal development have struggled to implement the code and adapt it to their unique studies. Furthermore, ADAE GUI was unable to process movies over 4.29GB which, given larger camera chips and faster imaging capacity, have become common and therefore must be addressed. Finally, batch processing of data sets proved problematic, impacting usability.

To build a more robust and versatile toolset for automated detection and unbiased analysis of exocytosis, we have developed a workflow in ImageTank (https://www.visualdatatools.com/ImageTank/, (O’Shaughnessy et al., 2019)), that we have called pHusion. ImageTank is a graphical programming language that combines image visualization with efficient numerical methods and an integrated interface to implement external code written in C^++^ (Stroustrup, 2013) or python (van Rossum, 1995). The data flow architecture employed by ImageTank enables visual inspection of each data transformation or calculation in real time, facilitating the identification of appropriate parameters and detection of points of failure. Further, to distinguish valid exocytic events, our previous analysis tools relied on particle tracking algorithms that can be opaque and unintuitive to adapt for different data sets. Instead of using these tools, we defined a set of relatively simple criteria such as permissible x-y drift and event duration that can be easily visualized and adjusted based on the experimental design.

Finally, as in previous work, we employ Ripley’s K-based analysis (Ripley, 1976; Ripley, 1977; Urbina et al., 2018) to determine if the spatial distribution of vesicle fusion events are clustered, uniform, or dispersed. Previously, our automated analysis made assumptions regarding the shape of the cellular or subcellular region to account for edge effects in the analysis (Urbina et al., 2018). Here, we present a fully generalizable approach in which no assumptions regarding cellular shape are made. Instead, we rely entirely on experimentally determined cell segmentations. This more general method is better suited to the disparate cell types presented in this work, and is thus more versatile. Further, here we have implemented temporal analysis based on Poisson processes to assess if exocytosis occurs in random bursts or is regulated in time.

To develop this more robust and flexible processing pipeline we have characterized exocytosis in diverse cell types including developing primary murine cortical neurons at two days in vitro (DIV), mature rat hippocampal neurons (DIV12-14), a human melanoma cell line (1205^Lu^), and primary rat oligodendrocytes. Further, we have used different pH-sensitive fluorophores (sepHluorin, pHmScarlet) tagged to various vSNAREs (VAMP2, VAMP3, VAMP7) imaged with Total Internal Reflection Fluorescence (TIRF) microscopy, Highly Inclined and Laminated Optical Sheet (HILO) microscopy, and widefield microscopy, and different camera sensor technologies. pHusion was adjusted for individual data sets and worked reliably in each scenario. We were able to capture markedly different frequencies of VAMP3-mediated exocytosis between oligodendrocytes and melanoma cells. Additionally, we demonstrate that the duration of constitutive VAMP2-mediated exocytic events in mature rat hippocampal neurons is much longer than that of immature murine cortical neurons.

## Results

To image exocytosis we employed a pH-sensitive variant of GFP, superecliptic-pHluorin (sepHluorin) fused to the vSNARE VAMP2. SepHluorin is quenched in the acidic lumen of the vesicle and fluoresces rapidly following fusion and exposure to the neutral pH of the extracellular environment (**Figure 1A**). After fusion, fluorescence decays exponentially as sepHluorin-VAMP2 either diffuses in the plasma membrane or is quenched following vesicle closure and reacidification (Bowser and Khakh, 2007; Urbina et al., 2021). Imaging VAMP2-sepHluorin in developing cortical neurons, we found that our initial analysis pipeline, ADAE GUI, performed within ImageJ, MATLAB, and R was brittle and failed frequently for a variety of reasons. To build a more robust and transparent analysis package, we used ImageTank for generalized image processing functions and custom code written in C^++^ for more specialized actions specific to exocytosis. All external code is launched from within ImageTank and results are passed back to it for further processing steps, thus avoiding the need to manually transfer data between separate applications. For our initial work to develop pHusion, we used immature primary cortical murine neurons expressing VAMP2-sepHluorin as a model system.

**Figure 1:**
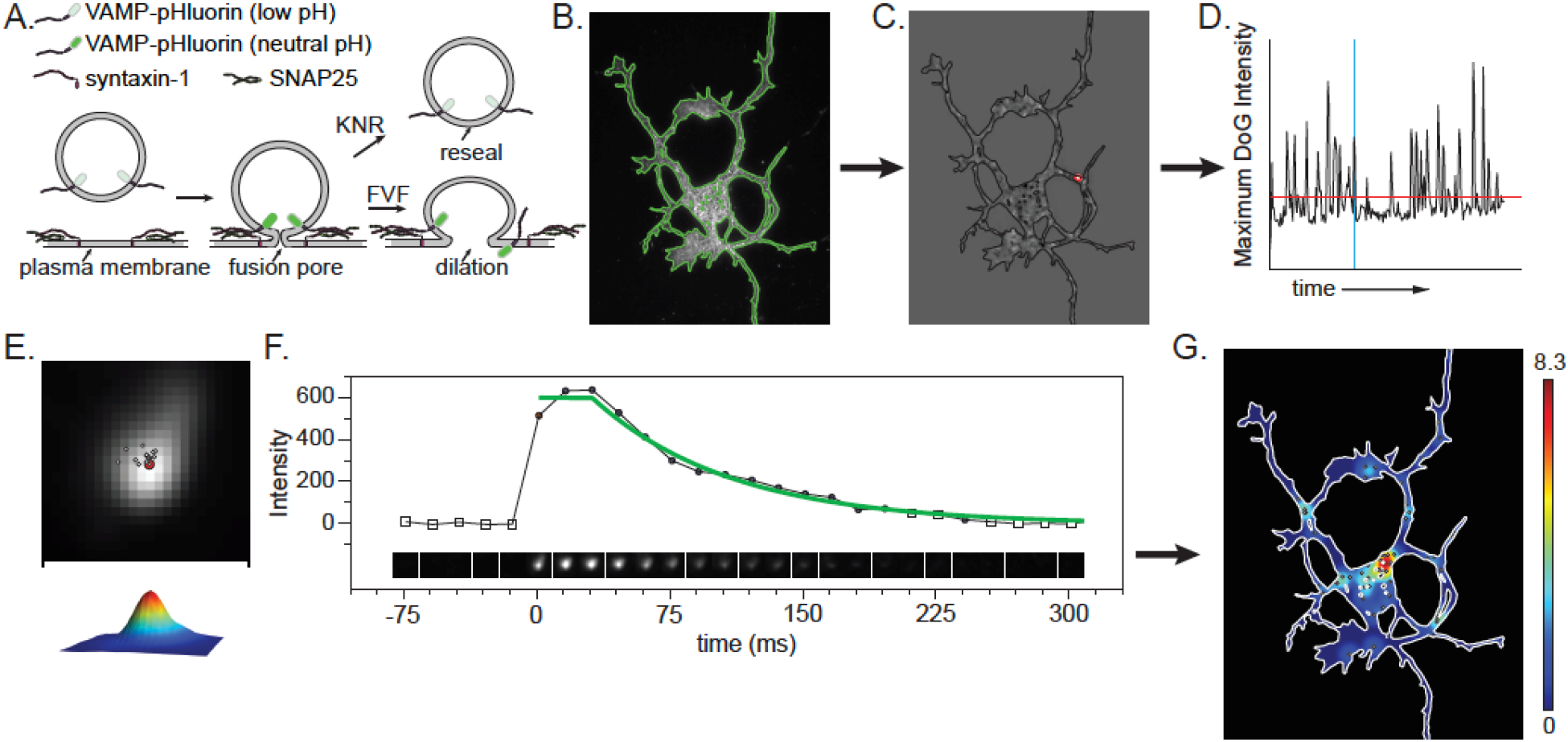
Schematic of pHusion workflow. **A.** Schematic of pH-sensitive sepHluorin fused to a vSNARE, with full vesicle fusion (FVF) and Kiss and Run (KNR) modes of fusion depicted. **B.** The images were segmented (green line) on the first frame and preprocessed as described in the text. **C.** Example difference of Gaussian (DoG) image with region of interest highlighted in red. **D.** The maximum intensity in the DoG image plotted over time. The threshold, shown as a red horizontal line is used to identify regions to evaluate further. The red contour shown in C is the region above the threshold for that time point. The vertical blue line identifies the time point displayed. **E.** A Gaussian model was fit to each cropped image and the center of the peak was used to quantify drift. Only those images with a goodness of fit above a defined value are included in the calculation, these timepoints are shown as circles in panel F. **F.** Example of an image series centered on a region of interest, the mean intensity in each image and the fit to the intensity values from the start of the event. **G.** Summary of all events detected and their spatial distribution.

### pHusion Workflow

We started with the basic framework for identifying potential exocytic events established in our previous work (Urbina et al., 2018). Briefly, the cell was segmented on the first frame, as the movies were short enough to ignore minor changes in shape (**Figure 1B**). The movie was preprocessed to subtract the background and photo bleach corrected by histogram matching to the fluorescence intensity of the first frame. Bright stationary objects were removed by subtracting a rolling window median projection of the preceding five frames. To filter out noise and enhance bright regions with the Gaussian profile of meaningful fluorescence, we used a difference of Gaussian (DoG) approach, subtracting successively blurred images and taking the median projection of the resulting images (**Figure 1C**). The maximum intensity in each projection was plotted over time and a threshold was applied to identify regions to investigate further (**Figure 1D**, red line). Though we followed the same basic steps as our previous work, a critical difference here is that we perform all calculations on the data as floating-point numbers, allowing decimals, not integer values. We chose to use floating-point numbers as maintaining only integers reduced the complexity of the data too much to enable fine-tuning the threshold parameter and was a frequent source of inaccurate identification (**Supplemental Figure 1**). Further, in the ADAE GUI images were reduced to 8-bit images to increase processing speed and reduce data size. Though useful, this further loss of data was unnecessary given the improved efficiency of calculations and flexible data caching approach used in pHusion.

For each region identified by the DoG, we cropped a small image series centered on the object in the raw image and extracted a defined number of frames before and after the event was detected. The size of the cropped image was 25×25 pixels for all of the data presented here, but is an adjustable parameter in pHusion. After applying a Gaussian filter of 1, a local background subtraction was performed using the median of the time points before detection and a Gaussian model was fit to each cropped image (**Figure 1E**). SepHluorin permits visualization of exocytic events due to the change in pH of the acidic vesicle lumen upon exposure to the extracellular environment, however vesicles that are not sufficiently acidified will maintain fluorescence of a distinct profile and must be ruled out from bona fide exocytic events. Instead of relying on particle tracking algorithms to identify exocytic events as we did in the ADAE-GUI, here we developed a simple set of rules to determine if an identified fluorescent event was a vesicle undergoing exocytosis. We defined exocytic events to be the transient appearance of Gaussian fluorescence above a threshold with minimal drift in x-y that undergoes exponential decay. These rules eliminate moving vesicles involved in trafficking and long lasting fluorescent puncta that do not undergo decay or diffuse fluorescent fluctuations. To accomplish this, each potential event was evaluated for the intensity above background, the drift, and duration. Further, a measure of intensity (either the maximum or mean) was fit with a two-step function consisting of a plateau followed by an exponential decay (**Figure 1F**). To determine the length of the plateau, we calculated the goodness of fit for the function starting with no plateau and increasing in length until end of the image series. The best fit was selected. We chose to make these parameters as explicit and accessible-as possible, allowing them to be easily adapted to different cell types and/or imaging conditions. With flexibility in mind, the code we have written in C^++^ enables modification of all key input parameters in ImageTank directly from the level of Gaussian blurring in the DoG, to the stringency of the drift calculation, and the goodness of fit of the exponential decay (**Table 1**). To facilitate optimization and quality control, we report metrics for every region analyzed and return flags to indicate the reason for exclusion such as low intensity, drift, or short duration. Finally, despite best efforts, automated analysis will inevitably include some failures and thus we have included a manual override to either include erroneously eliminated events or remove false calls. This manually curated list is then used for further analysis such as spatiotemporal analysis or classification. In its simplest form pHusion is performed on one timelapse movie at a time, though the analysis is amenable to batch processing, which can reduce the number of user interventions and greatly accelerate analysis time.

**Table 1:**
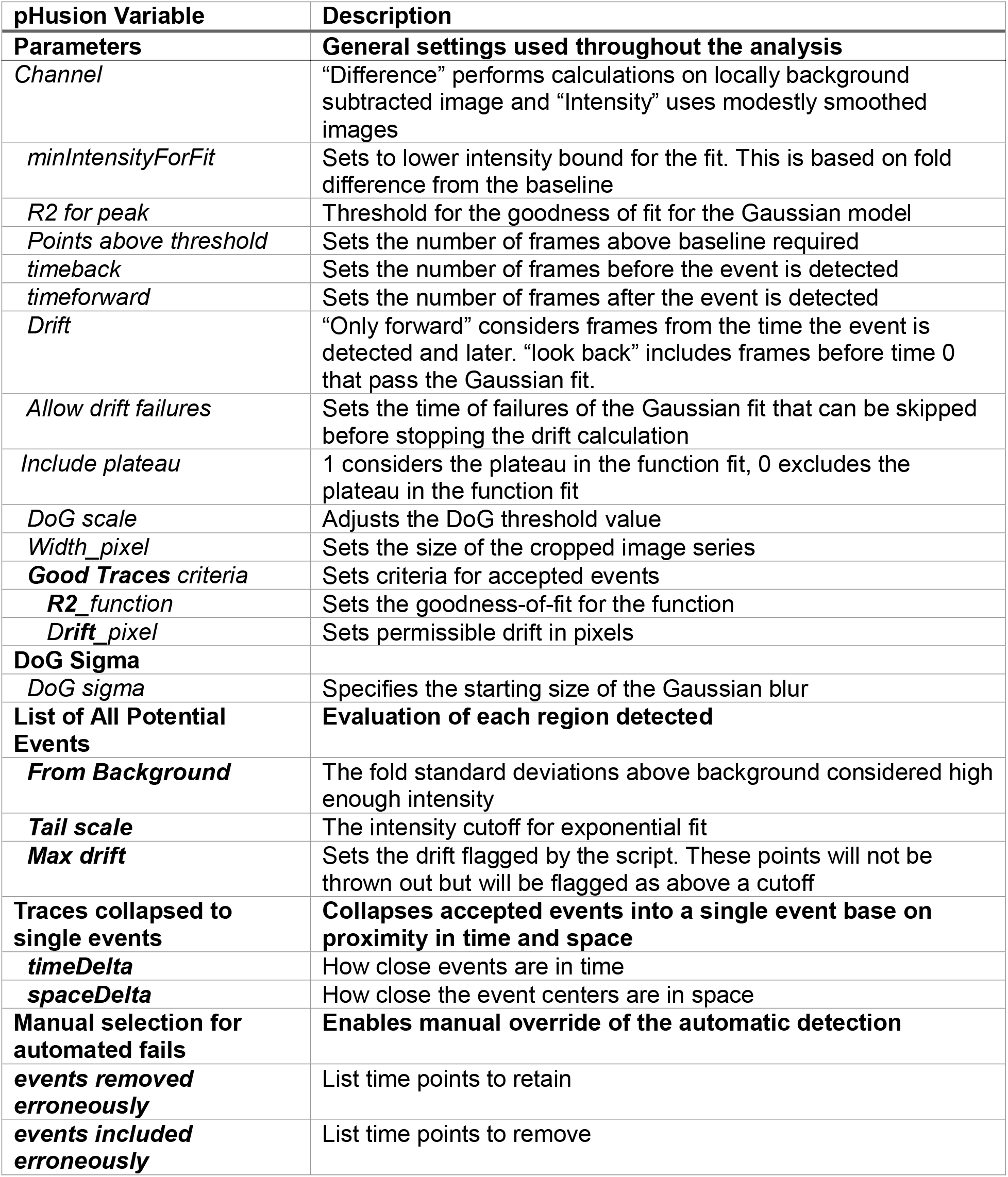
Parameters that can be adjusted to adapt the analysis for different cell types and/or imaging conditions. Specific sections of the script that can be modified are shown in bold. The individual options that can be adjusted in each section are listed below.

### Comparison of ADAE GUI and pHusion

We were motivated to develop this more robust analysis tool because we found that ADAE GUI failed frequently for new users, in different cell types, and/or for data sets acquired by distinct imaging paradigms. For example, when we changed our camera from an EMCCD to a high-speed sCMOS, the analysis failed completely for all movies over 4.29GB, as well as for unknown reasons on smaller datasets. For those cells for which results were returned, the set of frequencies calculated was far more varied with several clearly spurious results (**Figure 2A**, sCMOS GUI). Using pHusion we identified consistent and comparable exocytic frequencies in murine cortical neurons expressing VAMP2-sepHluorin that resulted in no failures regardless of camera type (**Figure 2A**, pHusion).

**Figure 2:**
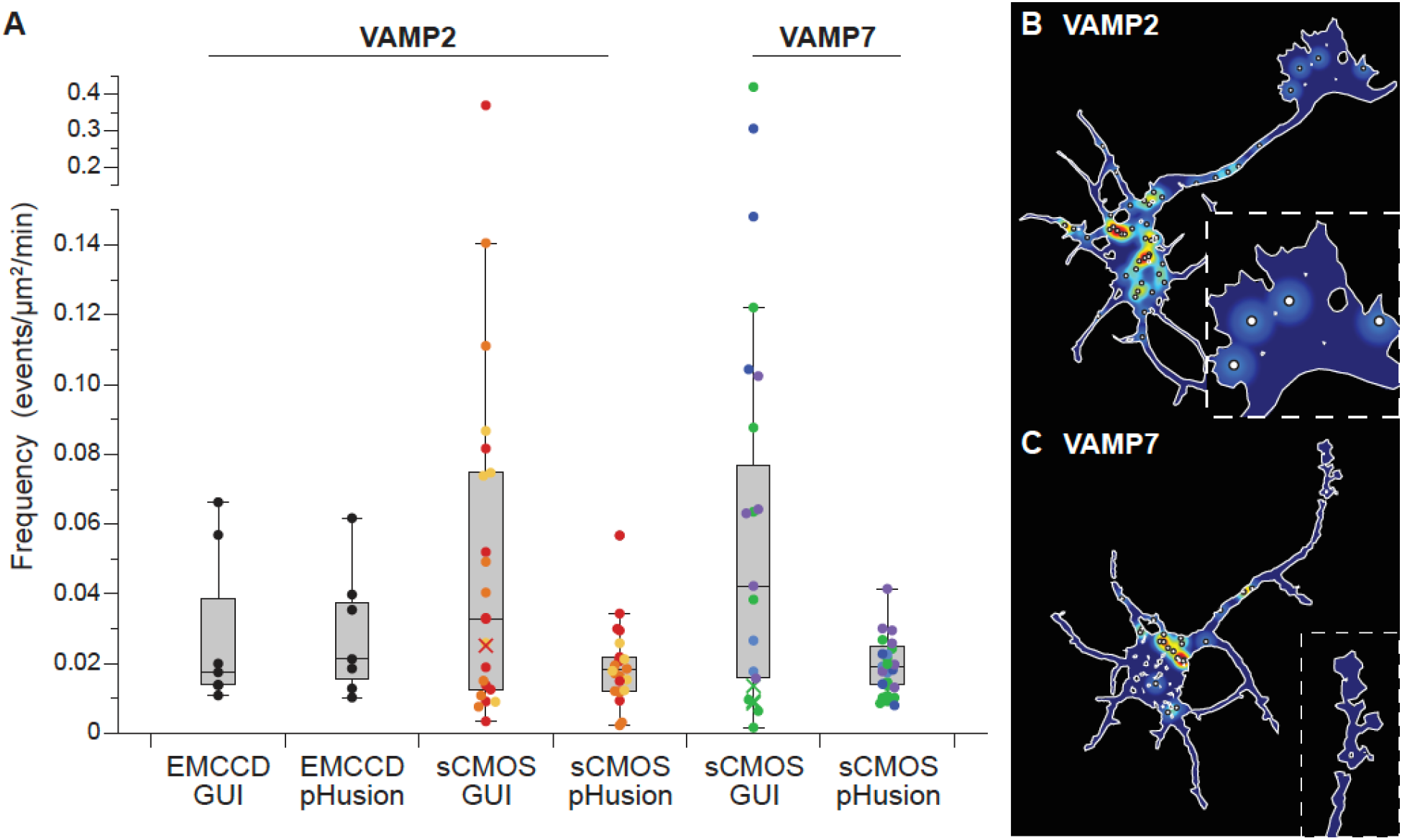
Comparison of ADAE GUI and pHusion analysis. **A.** Primary murine cortical neurons at 2 days in vitro (DIV) expressing VAMP2-sepHluorin were imaged with either an EMCCD (n = 7 cells) or sCMOS (n = 24 cells) camera and analyzed with both the ADAE GUI and pHusion to automatically detect exocytic events. The frequency of exocytosis reported as no. of events/µm^2^/min were: EMCCD-GUI: 0.029 ± 0.0087, EMCCD-pHusion: 0.029 ± 0.0070, sCMOS-GUI: 0.058 ± 0.017, sCMOS-pHusion: 0.024 ± 0.002. VAMP7-sepHluorin expressing cells (n=27) were imaged with a sCMOS camera and analyzed with pHusion (0.019 ± 0.0015) and the GUI (0.079 ± 0.021). **B-C.** Primary murine cortical neurons at 2 days in vitro (DIV) expressing VAMP2-sepHluorin (B) or VAMP7-sepHluorin (C) were imaged with sCMOS.

VAMP7 is another vSNARE enriched in developing neurons that has been implicated in exocytosis and neuronal morphogenesis (Coco et al., 1999; Gupton and Gertler, 2010; Martinez-Arca et al., 2001; Urbina and Gupton, 2020). Our previous work has shown discrepancies in the frequency of exocytosis mediated by VAMP7 in developing cortical neurons with rates similar to VAMP2 reported for manually identified events (Gupton and Gertler, 2010; Winkle et al., 2014) and considerably lower frequencies detected with ADAE GUI (Urbina et al., 2018). We sought to analyze VAMP7-sepHluorin imaging with pHusion. Compared to VAMP2-sepHluorin, VAMP7-sepHluorin expressing cells had a higher number of non-transient puncta and moving structures that confounded our analysis. This increase in fluorescent noise required us to adjust several parameters in the script to eliminate a great number of incorrectly identified events. To reduce the number of moving vesicles identified by pHusion, we tightened the permissible drift, included frames before the official start of the event provided they could be fit with a Gaussian model, and lowered the threshold for the goodness of fit for the model. Overall, these changes gave us more frames to track and improved our ability to remove these events. We also found that lowering the initial sigma value in the DoG reduced the number of diffuse puncta erroneously detected by the analysis. With these changes we reduced the error rate to under 10% and found the frequency of exocytosis to be similar to that of VAMP2-sepHluorin (**Figure 2A**, VAMP7). Further, consistent with our VAMP2-sepHluorin imaging, we found that analysis of VAMP7-sepHluorin expressing cells with ADAE GUI failed to produce results for a number of cells and, when results were obtained, the frequencies were much more varied and in some cases clearly inaccurate (**Figure 2A**, VAMP7 GUI). This provides a direct example of how the ability to adjust script parameters during data visualization benefits accurate analysis. We observed similar overall frequencies of exocytosis between VAMP2 and VAMP7. Mapping sites of exocytosis onto the neuronal mask also revealed more events occurring in the soma than the growth cone.

### Comparison of sepHluorin and pHmScarlet

Our initial studies of exocytosis were done with sepHluorin, a well-established and widely used fluorophore in the green spectral range. We wanted to test a newly developed red pH-sensitive fluorophore called pHmScarlet (Liu et al., 2021) as it was reported to be brighter than existing pH sensitive red proteins like pHuji(Shen et al., 2014) and can capture both docking and fusion events (Liu et al., 2021). We imaged primary cortical neurons at DIV2 expressing either VAMP2-sepHluorin or VAMP2-pHmScarlet and found a modest, but significant, reduction in detected exocytic frequency for pHmScarlet (**Figure 3A, Individual**). For a more direct comparison, we imaged both probes simultaneously using a Hamamatsu Gemini Image Splitter and saw an even greater decrease in detected frequency (**Figure 3A, Simultaneous**). This may be due in part to bleed-through correction required for the pHmScarlet channel when performing dual color imaging. We found that approximately 8% of the sepHluorin channel bled into the pHmScarlet image and therefore had to be corrected. However, we also detected significantly less events with each fluorophore expressed and imaged simultaneously when compared with the same reporter expressed and imaged individually (**Figure 3A**). Interestingly, we observed that some events were captured with both pHluorin and pHmScarlet (**Figure 3B)**, but a significant number of events were detected by only one reporter, either pHluorin (**Figure 3C**) or pHmScarlet (**Figure 3D**). We quantified all unique events detected with either reporter and found that the combined frequency was less than, but not significantly different, from VAMP2-pHluorin imaged individually. Our data suggest that expressing both VAMP2 reporters simultaneously hampers the ability to detect either one as efficiently as imaging them independently. Further, the frequency of exocytosis in cells expressing both VAMP2-pHluorin and -pHmScarlet is unlikely to be due to an overexpression artifact, as we found that the frequency of events does not depend on the expression level of VAMP2 (**Figure 3E**).

**Figure 3:**
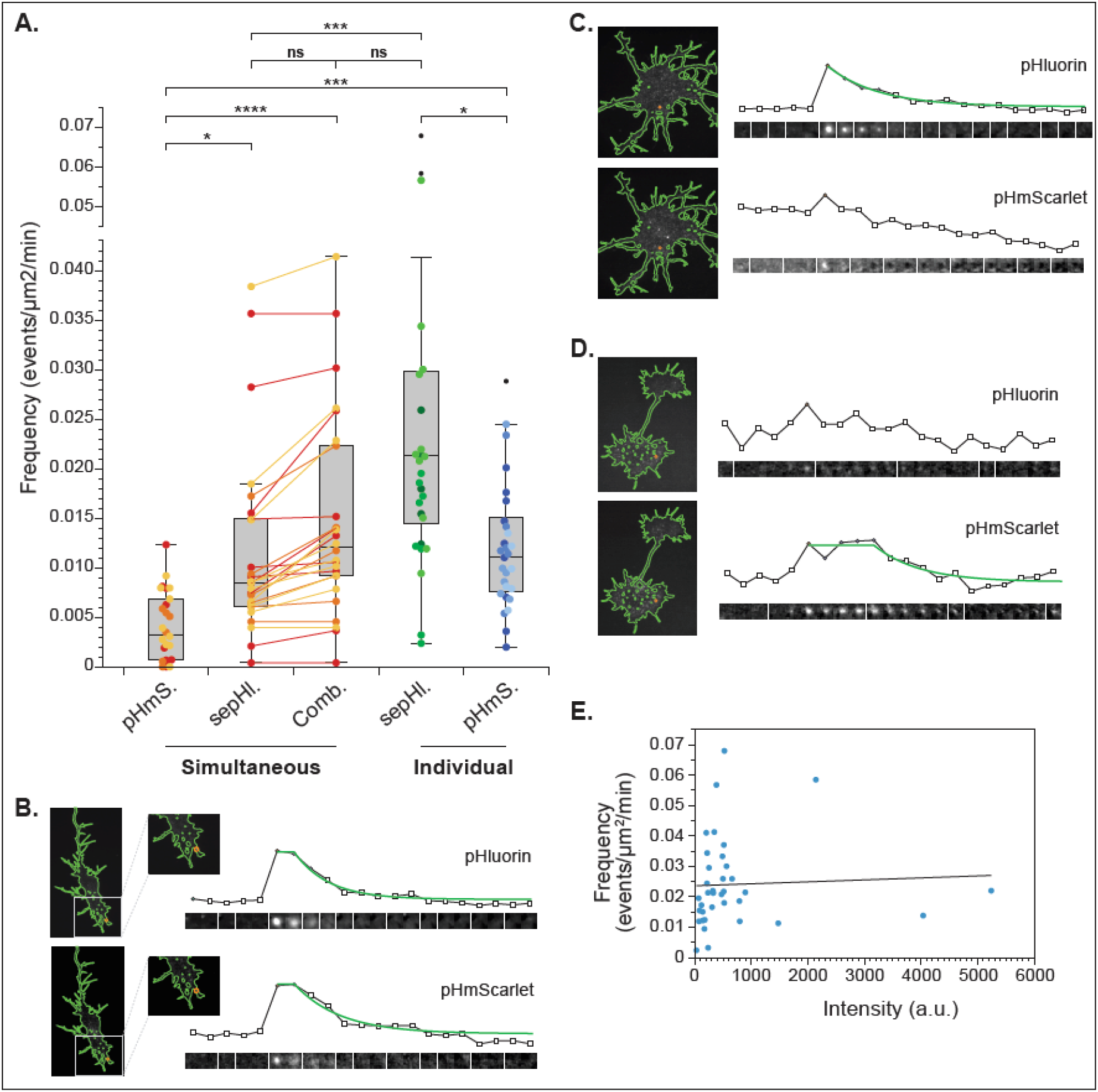
Comparison of VAMP2-sepHluorin and VAMP2-pHmScarlet in developing cortical neurons. **A.** Frequency of exocytosis in neurons expressing both VAMP2-sepHluorin and -pHmScarlet (n = 26) imaged simultaneously using a Hamamatsu Gemini Image Splitter and cells expressing VAMP2-sepHluorin alone (n=36) or VAMP2-pHmScarlet alone (n = 27) imaged separately. In simultaneous imaging the frequencies, reported as number of events/µm^2^/min, were pHmScarlet: 0.0041 ± 0.00068, pHluorin: 0.012 ± 0.0019, or combined frequency of unique events in either channel: 0.015 ± 0.0020. The frequency of VAMP2-pHluorin imaged separately was 0.024 ± 0.0024 and VAMP2-pHmScarlet was 0.012 ± 0.0012. **B.** Representative event observed with both pHluorin and pHmScarlet. **C.** Representative event observed with pHluorin but not pHmScarlet. **D.** Representative event observed with pHmScarlet but not pHluorin. **E.** Plot of frequency (number of events/µm^2^/min) against intensity (a. u.) for cortical neurons expressing VAMP2-sepHluorin alone. The linear function fit is y = 0.0235937 + 6.48894e-7*x with an R^2^ of 0.002.

### VAMP3-mediated fusion in oligodendrocytes and melanoma

To validate the versatility and flexibility of the exocytic detection further, we imaged primary rat oligodendrocytes at a mature stage of differentiation expressing VAMP3-sepHluorin using widefield epifluorescence microscopy. Mature oligodendrocytes are very large cells (∼120 µm in diameter) that require exocytosis during shape change (Lam et al., 2022) and are sufficiently flat to obviate the need for TIRF illumination. However the background fluorescence was higher than that observed by TIRF imaging and the intensity of exocytic bursts was much lower (**Figure 4A**). Surprisingly, despite these differences, we were able to adjust parameters in our script to successfully detect events. For oligodendrocytes, we increased the sigma for the DoG significantly and slightly lowered the stringency of the goodness of fit for the Gaussian model and the function fit. The frequency of exocytosis, though relatively low compared to cortical neurons at DIV2, was closely matched between automatic detection and manual analysis (**Figure 4B**). To confirm that the events identified were bona fide exocytic events, VAMP3-sepHluorin was co-expressed with mRuby-tagged tetanus toxin light chain (TeNT), which cleaves VAMP3 and blocks VAMP3-mediated SNARE complex formation (Galli et al., 1994). Exocytosis was quantified by both manual and automated detection. Regardless of the analysis method, the frequency of exocytosis was reduced to negligible levels by co-expression of mRuby-TeNT (**Figure 4B**), indicating identified events were bona fide exocytic events, and that the analysis pipeline in pHusion is sufficiently flexible for identification of exocytic events mediated by distinct vSNARES, in different cell types, imaged by widefield epifluorescence microscopy.

**Figure 4:**
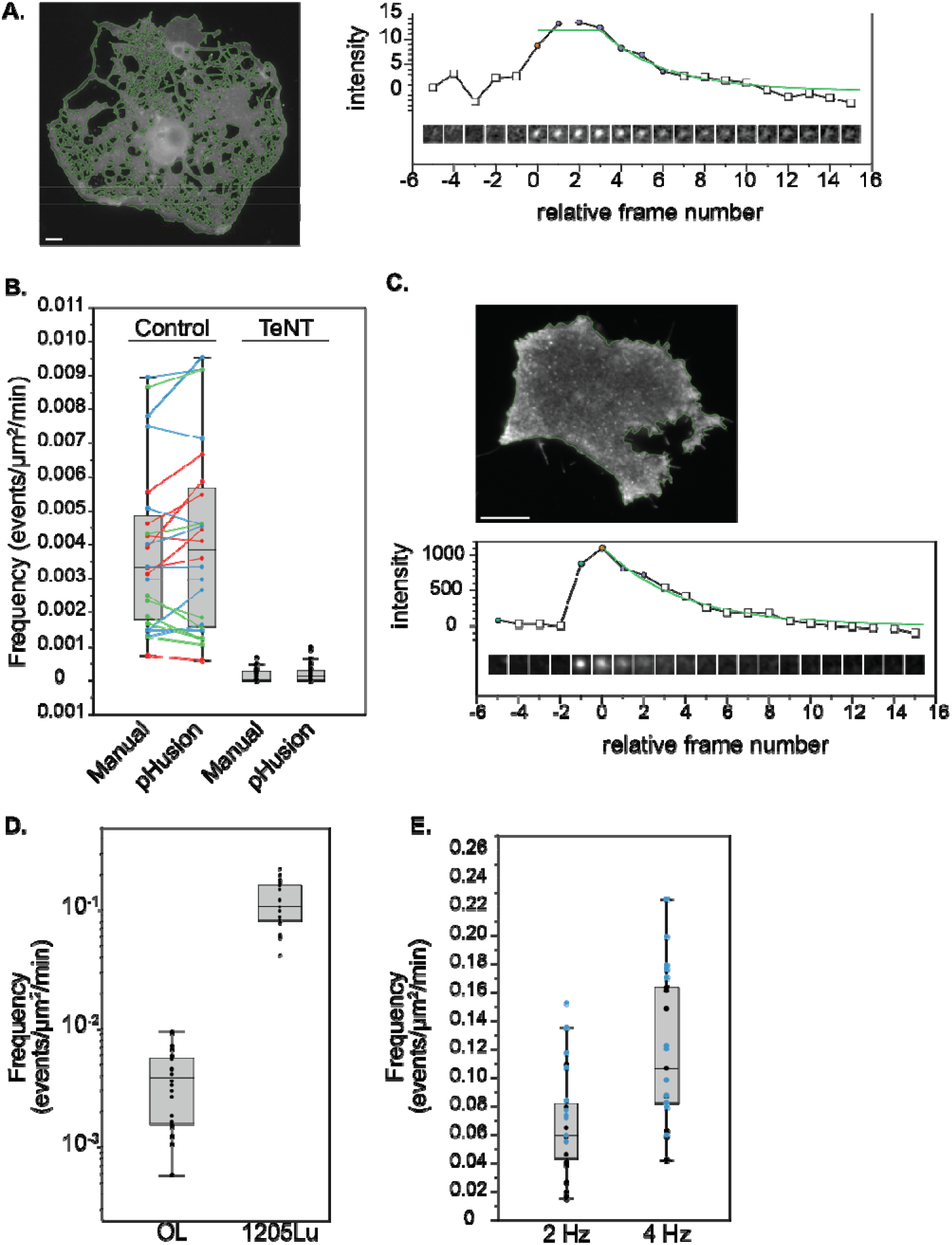
The frequency of VAMP3 exocytosis in primary rat oligodendrocytes is much lower than that of human melanoma cells. **A.** Representative image of segmented oligodendrocyte (OL) and event detail, scale bar 10 µm. **B.** Quantification of exocytosis with manual (0.0038 ± 0.0005) and automated (0.0041 ± 0.0006) detection (n=21 cells). Frequencies reported as number of events/µm^2^/min. Cells expressing both VAMP3-sepHluorin and tetanus toxin light chain (TeNT) were quantified manually (0.00012 ± 3.6e-5) and automated (0.0002 ± 5.6e-5). **C.** Representative image of 1205^Lu^ human melanoma cell line and example exocytic trace. **D.** Frequency of VAMP3-sepHluorin exocytosis in primary rat OL (0.0041 ± 0.0006) and 1205^Lu^ cells (0.12 ± 0.011, n = 21). **E.** Frequency of exocytosis in 1205^Lu^ cells imaged at 2 Hz (0.067 ± 0.0077, n = 23) and 4 Hz (0.12 ± 0.011, n = 21).

Secretion of matrix metalloproteases promotes metastasis of cancer cells. We next hoped to examine exocytosis in a metastatic cell line and determine how pHusion needed to be adapted for analysis of a mitotic, migratory cell type in which secretion is central to their pathology. We selected 1205^Lu^, a melanoma cell line with a metastatic phenotype (Simon et al., 1996). Interestingly, in contrast to the modest modification our analysis script required to process oligodendrocytes imaged with epifluorescence, images of VAMP3-sepHluorin in 1250^Lu^ cells acquired with the same TIRF imaging paradigm used above for cortical neurons, required significant alteration. A subset of exocytic events had a much smaller diameter than those in other cell types, and thus we had to lower the sigma in the DoG to prevent loss of these events in blurring. Further, the frequency of events was so high that the maximum intensity vector generated by the DoG did not have a clear demarcation between background and regions of interest. This required us to set a lower threshold for identifying regions for further evaluation in order to not miss events. However, lowering the threshold introduced fluorescent regions that were not exocytic events, and thus needed to be filtered. To do so, we tightened the stringency of the Gaussian fit and exponential decay to classify events as “true”. After making these adjustments, we achieved an error rate of less than 5% and found that the exocytic frequency of VAMP3 vesicles is far greater in human melanoma cells compared with primary rat oligodendrocytes (**Figure 4D**). The frequency of exocytosis we report is likely an underestimation, as we found that increasing the speed of acquisition increased the frequency obtained, but we could not image faster than 4 Hz due to phototoxicity (**Figure 4E**).

### Persistence of exocytic events in mature neurons

We also altered analysis parameters to enable automatic detection of exocytosis in mature primary rat hippocampal neurons (DIV 12-14). Segmenting these highly complex and varied cells proved to be time consuming and error prone and thus we performed the analysis on the entire image without masking the cell (**Figure 5A**). Interestingly, to image these fine, dim structures required camera settings that introduced shot noise detectable by the DoG analysis. These phenomena were exacerbated by the lack of a cell mask. Fortunately, shot noise is very rapid and does not move in x-y and thus could be eliminated based on duration and by requiring a minimum drift greater than zero. Further, we observed that many events were persistent with a much longer plateau than those observed in developing cortical murine neurons (**Figure 5B**). As a result, we had to lower the R^2^ threshold for the function fit. The R^2^ for a constant line is 0 and consequently inclusion of large segments of constant values artificially lowered the overall function fit.

**Figure 5:**
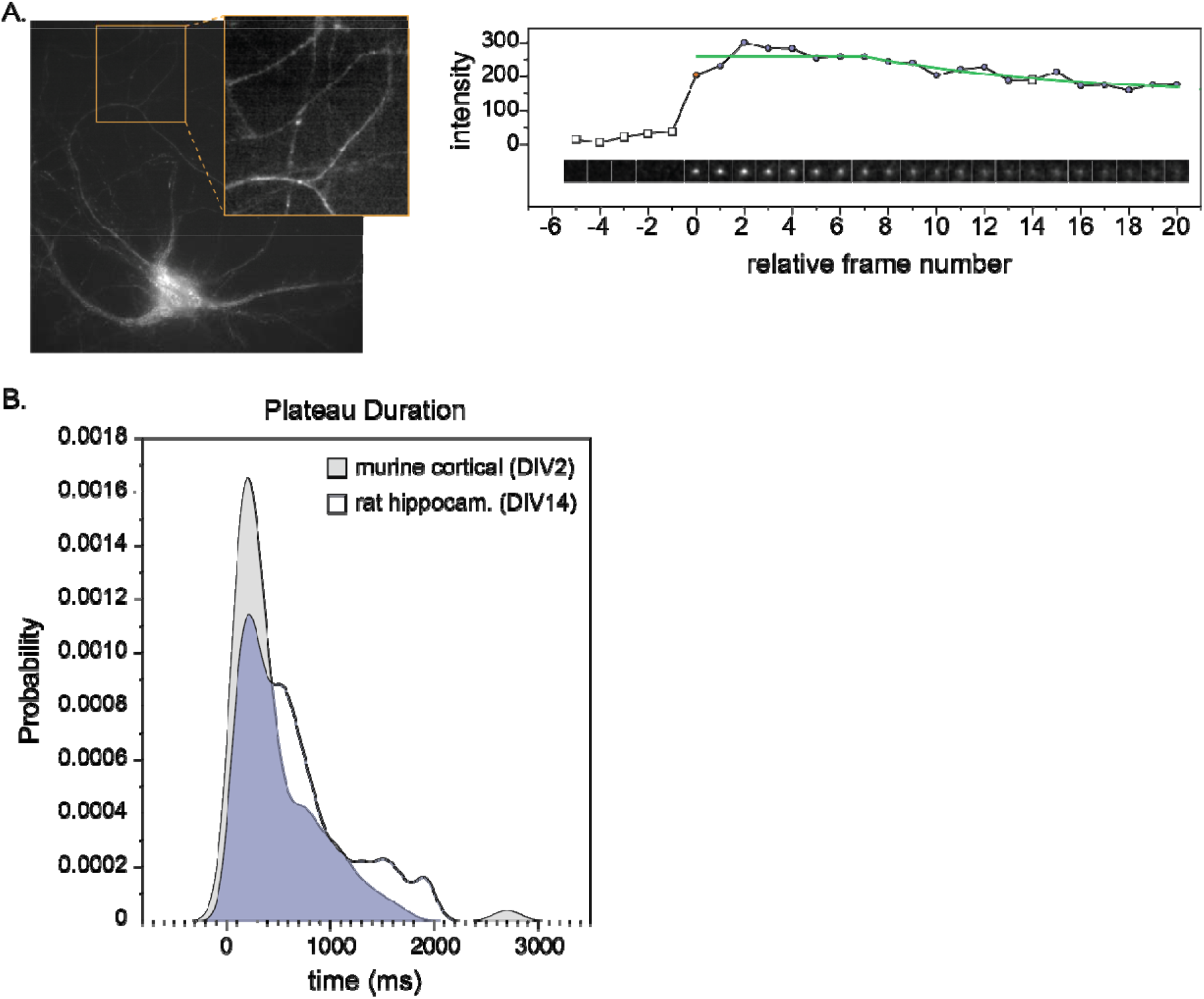
Persistence of exocytic events in developing and mature neurons. **A.** Representative image of primary rat hippocampal neurons (DIV 12-14) and a representative exocytosis trace. The inset shows a detail of the network. **B.** Histograms of the time of the event plateau between mature (n = 223) and immature neurons (n = 203).

### Spatio-temporal analysis exocytosis

Hotspots of vesicle fusion are frequently observed (**Figure 6A**), suggesting that the spatial distribution of exocytic events was not random. Mathematically, random events are defined as being distributed uniformly, whereas dispersed events are distributed in a regularly spaced arrangement. Discerning whether exocytic events are clustered, uniform, or dispersed requires spatial statistics. Our previous work to characterize the spatial distribution of exocytic events in developing neurons relied upon manual segmentation of the cell into the soma and neurites. We used Ripley’s K analysis to classify events as clustered, dispersed, or uniform (Urbina et al., 2018), and calculated an actual Ripley’s L(r). Because this analysis was developed for simple geometries, we used a weighting term to compensate for edge effects in the neurites, but not the soma.

**Figure 6:**
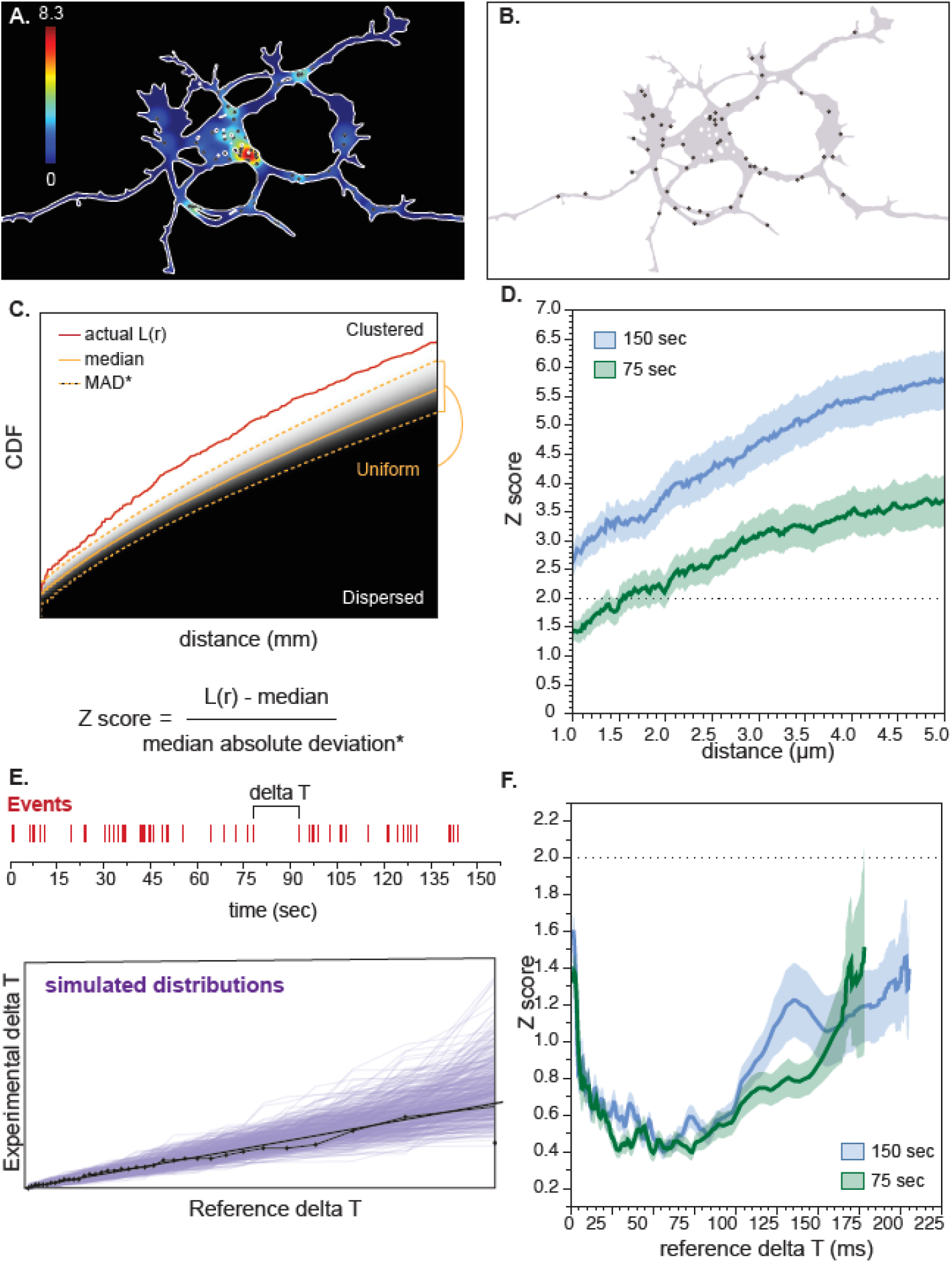
Generalized analysis of spatial distribution of exocytosis. **A.** Representative heatmap of an actual event distribution. **B.** Example of one Monte Carlo simulation of an event distribution using the same number of events detected and restricted to the cell mask. **C.** The overlay of the cumulative density functions for all 25000 simulations. The actual L(r) is shown in red, the median of the simulations is shown in orange and the bounds of the MAD are shown as orange dashed lines. An actual L(r) above the upper bound of the MAD is considered clustered, below the lower bound of the MAD is considered dispersed and between the bounds is taken to be uniform, or random. **D.** Spatial Z-score (mean ± SEM) for VAMP2-sepHluorin exocytosis in developing cortical neurons for full length, 150ms (n = 45) or half length, 75ms (n=36) movies. **E.**

Our goal in this work was to develop a generalized process to classify the spatial distribution of events that did not require manual input based on cell morphology. Using each cell mask and the number of events identified for that cell, we performed Monte Carlo simulations of events and plotted the cumulative distribution functions (CDF) for each of 25,000 simulations (**Figure 6B,C**). Using this collection of CDFs, we calculated the median and the non-parametric median absolute deviation (MAD) as a function of distance (r) to define the boundaries, above which events are clustered and below which, events are dispersed (**Figure 6C**). When distributed within the MAD, events are taken to be uniform, or randomly distributed in space. The actual Ripley’s L(r) for each cell was compared to its respective simulations. To enable comparison of a collection of cells regardless of differences in cell shape and the resulting CDFs, we calculated a non-parametric Z score as a function of r by taking the difference between the actual L(r) and the median divided by the MAD. Z scores above 2 (or below −2) are considered significant and are thus, not uniform. Consistent with previous work, we found that VAMP2-mediated events were clustered in space in developing cortical neurons at DIV2 (**Figure 6D**). Interestingly, though we explicitly exclude time from the analysis we do find that the duration of imaging does affect our results. For example, when we analyzed our full datasets acquired for approximately 2.5 minutes, we observed spatial clusters over the entire range of distances considered. However, when we analyzed just the first half of the movies (∼1 min), we no longer observed spatial clustering of exocytosis over short distances (< 1.5 µm), likely because we had not allowed sufficient time for repeated events to occur frequently enough in discrete hotspots.

In addition to measuring the spatial distribution of events, we wanted a robust method to determine if exocytic events were random in time. Random events in time can be described as a Poisson process, in which every event is independent of all others. By definition for a Poisson process measured over a given period of observation, the difference in time between all consecutive events will follow an exponential distribution. Thus, to determine if exocytic events were random in time, we measured the time step (delta T) between consecutive pairs of events (**Figure 6E**) and determined whether the set of delta T was exponentially distributed. To measure if a set of values was exponentially distributed, we used a quantile-quantile (Q-Q) plot of the experimental dataset against the standard exponential function with a decay rate of 1 (**Figure 6E)**.

When two datasets are drawn from the same underlying type of distribution, the Q-Q plot results in a line. Thus, we fit a line to the Q-Q plot and simulated 200 exponential distributions consistent with the fit line. A simple linear regression is not robust enough to experimental noise to rigorously determine if the data were consistent with an exponential distribution. Instead, the simulated distributions were used to generate an envelope against which experimental data can be tested. As with space, we calculated a Z score across time by subtracting the fit line from the actual data and divided by the MAD. Z scores above 2 were considered significant. To pool data from different cells each Z score was interpolated onto a common time axis. We found that exocytosis occurred as random bursts in time regardless of the overall duration of the movie (**Figure 6F**).

## Discussion

Due to the complexity and volume of image-based datasets generated by modern microscopy systems, computer-vision based approaches for analysis are warranted and, in some cases, essential. Though valuable, designing automated methods for quantitation of visual data is very challenging and often results in brittle methods that work only for specific data sets. This is the problem we faced with automated analysis of exocytosis using ADAE GUI. Our initial processing scheme worked very well for the data on which it was developed, but failed frequently when used to analyze even slightly divergent datasets.

To develop a more versatile and robust application we used ImageTank, software that seamlessly merges numerical methods and visualization of all data types from 3D image stacks to 1D data tables. The heavy reliance on visualization allowed us to rapidly detect key differences between datasets and identify ways to adapt the code for different datasets. Additionally, visualization provided built in quality control mechanisms as the user can quickly see all aspects of workflow from the threshold applied to the DoG to each individual event evaluated. The ability to adapt analysis to different imaging conditions and consequently generate reliable annotated datasets can be useful in developing deep learning approaches.

Further, improvements in camera technology both in terms of the speed of acquisition and the size of the chip has greatly enhanced resolution, but comes at the cost of the ever increasing size of image files. To accommodate larger file sizes, we use a hybrid memory/disk based and “just-in-time” computing approach that easily handles datasets too large to be stored in memory alone. With this data handling architecture, our script does not preclude analysis of large files well over the 4.29GB cutoff for so-called “big-tiff” files. Moreover, where data are stored, on disk or in memory, is actively managed to increase efficiency and calculations are heavily threaded to greatly improve overall processing sped. One limitation to our reliance on ImageTank is that the graphics, user interface, and threading rely heavily on the Mac operating system. Therefore pHusion is unavailable for other operating systems such as Windows or Linux.

The method we have developed to identify exocytic events depends heavily on the DoG to highlight transient Gaussian fluorescence that rises above a baseline. In some cases, such as the 1205^Lu^ cells, events occurred so frequently that a reliable baseline could not be determined. In this case a much lower threshold had to be applied and care taken to filter out erroneously identified regions. Another difficulty with the DoG method can arise when very small cells or structures are analyzed. Images need to have sufficient pixels to establish a baseline above the noise. For example, to visualize exocytosis in the growth cone we performed the analysis on the whole cell and then segmented the growth cone instead of analyzing the growth cone by itself as these data were too noisy and events too rare to accurately capture.

In addition to altering the parameters of our processing script to accommodate diverse datasets, adjusting the script for distinct imaging conditions was helpful and/or necessary as well. For example, though oligodendrocytes are sufficiently flat to enable epifluorescence imaging, the background was higher than in TIRF datasets. To address this, we photobleached VAMP3-sepHluorin within the PM prior to imaging exocytosis; we found this increase in signal to noise to be beneficial in imaging mature hippocampal neurons as well. While useful, care must be taken to ensure cells are not damaged by phototoxicity as sensitivity to light varies greatly across cell types. For example, we were unable to image 1205^Lu^ melanoma cells as rapidly as neurons, due to the deleterious effects of light.

We were interested in exploring a newly reported pH-sensitive fluorophore in the red spectral range, pHmScarlet, as this tool may enable us to distinguish docking and fusion events in exocytosis. Further, the flexibility of multicolor imaging experiments is greatly enhanced by the availability of tools of different wavelengths. We found that we detected a lower number of events using VAMP2-pHmScarlet compared with VAMP2-sepHluorin when these probes were imaged separately, and an even greater decrease was seen when imaged simultaneously. However, due to bleed-through from sepHluorin into pHmScarlet, we removed 8% of the sepHluorin image from pHmScarlet, which lowered the overall intensity in that channel. Nevertheless, taken together our data suggest that pHmScarlet may under estimate the frequency of exocytosis, at least with our imaging setup, but captures a subset of events.

Here we presented a generalized approach to assess the spatial distribution of exocytic events that made no assumptions regarding cell shape. Additionally, we presented a robust method to determine if events were random in time. These are useful tools in dissecting when and where exocytosis occurs and how it is regulated, but they are only a first step. Here, we found events to be random in time. For circumstances in which events are not random in time, further work is required to dissect whether these events exhibit periodicity or non-random bursting. Further, we have treated space and time independently. A more sophisticated approach would integrate space and time for a more complete picture of how exocytosis is regulated, but is beyond the scope of the current work.

## Materials and Methods

### ImageTank Analysis Script

#### Difference of Gaussian

For the difference of Gaussian (DoG), images were blurred with increasingly large Gaussian filters starting with a sigma of 3 for cortical neurons, VAMP2, 1 for VAMP7, 1 for 1205Lu melanoma, 8 for oligodendrcytes and 3 for hippocampal neurons. An image stack based on 13 rounds (k) of blurring was generated by subtracting image k+1 from image k and the median value of the image stack was calculated. The maximum intensity value in the median image was found for each time point. The median value of the maximum intensity vector was used to set the threshold above which identified regions (ROI) for further evaluation. The threshold was adjusted by multiplying the median value by a scale factor. For cortical neurons and oligodendrocytes, the value was scaled by 1.2, in hippocampal neurons by 1.5, and in 1205Lu by 0.7.

#### Evaluation of potential events

Using the ROI from the DoG, a small image series was cropped from the raw image stack taking a specified number of time frames before and after the spot was detected (defined as t = 0). These smaller images were smoothed with a Gaussian filter of 1 and were either used directly for subsequent analysis (channel “intensity” in the parameters list) or were background subtracted (channel “difference” in the parameters list). For background subtraction, the median of the images preceding t=0 was subtracted from the entire image series. The image just before the start of the event (t=-1) was excluded from the median as this often has elevated fluorescence intensity. Each image in the series was fit with a Gaussian model. The R^2^ (goodness-of-fit) for the Gaussian model was set to 0.6 for cortical neurons, 0.5 for oligodendrocytes, 0.75 for 1205Lu, 0.3 for hippocampal neurons. To be a valid event, the image at the time point the event was detected must be above the R^2^ for the Gaussian model and only frames above this threshold were considered in the drift calculation. The centers of the Gaussian models were used to calculate the maximum distance between all valid frames. A measure of the intensity (maximum or average) was fit with a two-step function consisting of a constant plateau and an exponential decay. The R^2^ for the function fit was 0.24 for hippocampal neurons, 0.7 for cortical neurons, 0.75 for 1205Lu and 0.5 for OLs.

#### Spatial analysis

Monte Carlo simulations using the number of events detected for a given cell and restricted to the mask were performed 25000 times. The collective cumulative distribution functions of these simulations were used to find the median probability of events as a function of radius (r) for the specific geometry of the cell. The actual L(r) determined from real events was compared to the median as follows:

Z-score = (L(r) – median) / (median absolute deviation (MAD)*)

MAD* = median(| xi – median(xi) |) / Φ^-1^*(0.75)

#### Temporal analysis

The time between sequential pairs of events were calculated for each cell to generate the delta T set. To assess whether the set of delta T values followed an exponential distribution used a quantile-quantile (Q-Q) plot. For each cell, we sorted the delta T and assigned percentile values as follows:

p = Entry # / (number of events + 1)

The actual delta T were plotted against theoretical delta T drawn from the standard exponential function with λ = 1, f(t) = e^-t^. The theoretical delta T are calculated at the p values as follows:

delta T = -ln(1-p)

A variance weighted line was fit to the resulting Q-Q plot and 400 exponential distributions were simulated that were consistent with the fit line to generate the Q-Q envelope. To compare the experimental data to the Q-Q envelope we used a Z-score similar to that calculated in space.

Z-score = (actual data point – fit line) / MAD*

To collect all cells into an aggregate Z-score each individual Z-score was interpolated onto a standard time scale and the mean and SEM were calculated for each time point.

Cells were excluded from both spatial and temporal analysis if fewer than 26 events occurred as the simulations were too noisy to be meaningful.

### Primary Murine Cortical Neuron Methods

#### Media, culture, transduction and transfection of primary murine cortical neurons

All mouse procedures were approved by University of North Carolina’s Institutional Anima Care and Use Committee (IACUC) and followed the National Institutes of Health guidelines. C57Bl6 male and female embryos were used for primary cortical neuron preparations.

Cortical neuron cultures were prepared from day 15.5 embryos as previously described (Viesselmann et al., 2011). Briefly, cortices were dissected and neurons were dissociated with trypsin and plated on poly-d-lysine–coated glass-bottom culture dishes in neurobasal media supplemented with B27 (Invitrogen). This same media was used for all time-lapse experiments. For all cells imaged with the cMos camera, VAMP2-SEpHluorin expression was mediated by adenoviral transduction, whereas for the EMCCD expression was mediated by nucleofection. Briefly, neurons were resuspended after dissociation in solution (Amaxa Nucleofector; Lonza) and electroporated with a nucleofector according to manufacturer protocol. For all other experiments exogenous expression was mediated by adenoviral transduction. Briefly, 36-48 hrs before imaging 2-5 ml of concentrated virus was added to each dish of plated cells.

#### Adenovirus production

Adenovirus constructs were produced following the manufacturer’s instructions (Takarabio Inc., Adeno-X™ Adenoviral System 3 (Tet-On® 3G Inducible)). Briefly, Adeno-X 293 cells (Takarabio Cat #632271) were transfected with linearized adenoviral vectors using Lipofectamine and Plus Reagent (Thermo Fisher). Cells were dislodged with gentle agitation upon exhibiting late cytopathic effect and were lysed with three freeze-thaw cycles. Pooled cell lysate and the flask supernatant was applied to a fresh flask of Adeno-X 293 cell for two rounds of amplification. Cell lysate alone was harvested by freeze-thaw cycles in PBS (Fisher Scientific, #MT-21-031-CV) following the final round of amplification. Expression of the target protein was induced with 1Lg/ml doxycycline.

#### Time-lapse Imaging

All TIRF imaging was performed using an inverted microscope (IX83-ZDC2; Evident/Olympus) with Cellsens software (Evident/Olympus), a cMOS camera (Orca-Fusion, Hamamatsu) or an electron-multiplying charge-coupled device (iXon) where indicated, and a live cell imaging chamber (Tokai Hit Stage Top Incubator INUG2-FSX) maintained at 37°C, CO2 at 5% and humidified. DIV2 neurons in culture media expressing VAMP2-sepHluorin, VAMP2-pHmScarlet and/or VAMP7-sepHluorin were imaged at 10 or 20 Hz with a 100× 1.50 NA TIRF objective and a solid-state 491-nm laser illumination with 30% power and a 110-nm penetration depth and/or a solid-state 561-nm laser illumination at 30% power at 110-nm penetration depth. Simultaneous dual-color image acquisition was performed with the Hamamatsu W-VIEW GEMINI image splitter optic (A12801-01).

#### Statistical analysis

For two-sample comparisons of normally distributed data, an unpaired t test was used for two independent samples, or a paired t test for paired samples. For non-parametric data we used the Kolmogorov-Smirnov (K-S) test to compare two-samples and the Kruskal-Wallis test for ANOVA. Spatial and temporal analyses were tested with a non-parametric Z-score with a cut off of 2.

### Primary Rat Oligodendrocyte Methods

#### Purification and culturing of primary oligodendrocytes

All rodent procedures were approved by Stanford University’s Administrative Panel on Laboratory Animal Care (APLAC) and followed the National Institutes of Health guidelines. Sprague-Dawley rats were ordered from Charles River Laboratories. Both male and female rat pups were used for primary oligodendrocyte culture preparations.

Primary oligodendrocyte precursors were purified by immunopanning from P5-P7 Sprague Dawley rat and P6-P7 transgenic mouse brains as previously described (Dugas and Emery, 2013; Emery and Dugas, 2013). Oligodendrocyte precursors were initially seeded on 10-cm dishes coated with 0.01 mg/ml poly-D-lysine hydrobromide (PDL, Sigma P6407) at a density of 150,000-250,000 cells. Cells were allowed to recover for 4 days in proliferation media containing serum-free defined media (DMEM-SATO base medium) supplemented with 4.2 μg/ml forskolin (Sigma-Aldrich, Cat#F6886), 10 ng/ml PDGF (Peprotech, Cat#100-13A), 10 ng/ml CNTF (Peprotech, Cat#450-02), and 1 ng/ml neurotrophin-3 (NT-3; Peprotech, Cat#450-03) at 37°C with 10% CO_2_. (Dugas and Emery, 2013; Emery and Dugas, 2013).

#### Transfection of oligodendrocyte precursors

Proliferating rat oligodendrocyte precursors were lifted from tissue culture dishes, and centrifuged at 90xg for 10 min. 250,000 oligodendrocyte precursors were gently resuspended into 20 ml of nucleofector solution (Lonza P3 Primary Cell 4D-Nucleofector V4XP-3032) with 300 ng of VAMP3-pHluorin and 300 ng of mRuby-caax or TeNT-mRuby-caax plasmids. Cells were then electroporated in a Lonza 4D-Nucleofector X Unit (AAF-1003X) assembled with a 4D-Nucleofector Core Unit (AAF-1002B) with pulse code DC-218. Electroporated cells rested for 10 min at RT before resuspension in antibiotic-free DMEM-SATO base media supplemented with differentiation factors, containing 4.2 μg/ml forskolin (Sigma-Aldrich, Cat#F6886), 10 ng/ml CNTF (Peprotech, Cat#450-02), and 40 ng/ml thyroid hormone (T3; Sigma-Aldrich, Cat#T6397).

Each batch of 250,000 cells was distributed into 4 35-mm dishes with No. 1.5 glass coverslips (MatTek Corporation P35G-1.5-20-C) coated with PDL-borate (0.01 mg/ml PDL, which was first resuspended at 100x in 150 mM boric acid pH 8.4) for an optimal cell density. Each dish was half-fed with freshly supplemented DMEM-SATO media every two days. After 5 days, media was replaced with FluoroBrite DMEM-SATO (made with Fisher Scientific A1896701) supplemented with differentiation factors for 2 hours at 37°C, 10% CO_2_ before imaging.

#### Live-cell imaging of exocytosis in primary oligodendrocyte-lineage cells

Time-lapse imaging of exocytic events was performed on a Zeiss Axio Observer Z1 inverted microscope equipped with a Zeiss Axiocam 506 monochrome 6-megapixel camera, a stage top incubator (Okolab, H301-K-Frame) set to 37°C, and a digital gas blender (Okolab, CO2-UNIT-3L) set to 10% CO_2_ during image acquisition. Samples were imaged with a Plan-Apo 63x/1.40 Oil objective using widefield epifluorescence with a 12V halogen lamp. Due to a high baseline level of VAMP3-pHluorin intensity on the cell surface, cells were subjected to initial “pre-bleaching” consisting of 20 50-ms exposure at a 250-ms frame rate. Then, time-lapse sequences for exocytotic events were captured using an acquisition rate of 250 ms/frame for 1 min using the Zen Blue software. Images were viewed using Fiji/ImageJ software.

#### Cloning of plasmids used for oligodendrocyte transfections (VAMP3-pHluorin, mRuby-caax, TeNT-mRuby-caax)

For live imaging of exocytosis in cultured oligodendrocyte precursors/oligodendrocytes, the VAMP3-pHluorin reporter consists of rat VAMP3 conjugated to super ecliptic pHluorin (referred to as pHluorin) cloned into a pAAV vector backbone with a CMV promoter. DNA containing mRuby and tetanus toxin light chain (TeNT) were gifted from the Stanford Gene Vector and Virus Core and were cloned into a pAAV vector backbone with a myelin basic protein promoter (pMBP)(Gow et al., 1992). The pMBP drives expression of mRuby/TeNT at later stages (Day 3+) of oligodendrocyte differentiation.

### Mature Rat Hippocampal Neuron Methods

#### Neuronal culture

Primary neuronal cell culture was obtained in a similar procedure as previously described (Bingham et al., 2022). Briefly, hippocampi were extracted from E18 rat pups from pregnant female Wistar rats (Janvier labs), dissected, and homogenized by trypsin treatment followed by mechanical trituration and seeded on 18-mm diameter round, #1.5H coverslips at a density of 30,000 cells/cm2 for 3 hours in serum-containing plating medium (MEM with 10% fetal bovine serum, 0.6% added glucose, 0.08 mg/mL sodium pyruvate, 100 UI/mL penicillin-streptomycin). In accordance with the Banker method (Kaech and Banker, 2006), the coverslips (cells facing down) were then transferred to and cultured in petri dishes containing confluent glia cultures conditioned in NB+ medium (Neurobasal medium supplemented with 2 % B-27, 100 UI/mL penicillin/ streptomycin and 2.5 µg/mL amphotericin).

All procedures were in agreement with the guidelines established by the European Animal Care and Use Committee (86/609/CEE) and was approved by the local French ethics committee (agreement G13O555).

#### Neuronal transfection and HILO microscopy

Neurons were transfected with 1 μg of VAMP2-phluorin between 12 - 14 DIV with Lipofectamine 2000 (Thermo Fischer). After 24 hours, live-cell imaging of neurons expressing VAMP2-phluorin was performed on an inverted microscope ECLIPSE Ti2-E (Nikon Instruments) equipped with an ORCA-Fusion sCMOS camera (Hamamatsu Photonics K.K. - C14440-20UP) and a CFI SR HP Apochromat TIRF 100X Oil (NA 1.49) objective. The system was equipped with a Nikon Perfect Focus System (PFS) and images were acquired with the NIS-Elements AR 5.30.05 software. Coverslips with neurons were mounted in a metal chamber in Tyrode’s solution (in mM: 119 NaCl, 25 HEPES, 2.5 KCl, 2 CaCl2, 2 MgCl2, 30 glucose, pH 7.4). Neurons were maintained in a humid chamber at 35.5°C-37°C for the duration of the experiments using a stage-top incubator (Okolab inc). To image exocytosis, VAMP2-SEP present initially in the plasma membrane was photobleached by exposing it to high power 488 nm laser light for 30 seconds before time-lapse imaging to remove the basal membrane signal and highlight exocytic signal (Yudowski et al., 2007). 100 ms exposure frames were then continuously acquired for 90s using 488 nm laser light at lower power (1-10 %).

#### Video preprocessing

Acquired videos were preprocessed using the *Filter Timelapse* script (available at https://sites.imagej.net/Christopheleterrier/plugins/NeuroCyto%20Lab/Kymographs/) that performed image stabilization using the Image Stabilizer plugin (https://imagej.net/plugins/image-stabilizer) after 2X downscaling, and bleach correction using average intensity compensation for the foreground identified as the top 12% of pixel intensities within each image of the sequence.

### Human Melanoma Cell Line Methods

#### Cell culture

1205Lucells were grown in DMEM (11995065;Gibco Thermo Fisher Scientific) supplemented with 10% FBS and 1% penicillin/1% streptomycin, at 37°C, in a humified environment with 5% CO2

#### Transfection and Imaging

1205Lu cells were plated on 35mm glass bottom dishes with 20mm micro-well #1.5 cover glass (D-35-20-1.5-n; Cellvis). For transient transfection,1205Lu cells were plated and 24 hours later were transfected with a plasmid encoding VAMP3-pHlourin using Lipofectamine (11668027; Life Technologies Inc) following the manufacturer’s instructions. After 16-18 hours, cells were imaged with a 100x 1.49 NA TIRF objective and a solid-state 491-nm laser illumination at 35% power at 100-nm penetration depth. Images were acquired every 500ms or 250 ms for 2 min with an exposure time of 50.995 ms.

## Acknowledgments

We would like to thank Prof. Steve Marron for his guidance on statistical measures for spatio-temporal analysis. We acknowledge funding from NIH R01NS112326 and 3R01NS112326-03S1 (SLG), C.L. would like to acknowledge funding by the Agence Nationale pour la Recherche (ANR-20-CE13-0024 to C.L.), and to thank the Neuro-Cellular Imaging Service and Nikon Center for Neuro-NanoImaging at INP, funded by CPER-FEDER (PlateForme NeuroTimone PA0014842). SER was supported by the Cellular Systems and Integrative Physiology Training Program T32GM133364. KL was supported by HHMI Gilliam Fellowship GT15781. JBZ would like to acknowledge funding from the McKnight Endowment Fund for Neuroscience (JBZ), the Stanford Bio-X Interdisciplinary Initiatives Seed Grants Program (IIP) [R9-24] (JBZ), the National Multiple Sclerosis Society Harry Weaver Neuroscience Scholar Award (JBZ), the Beckman Young Investigator Award (JBZ), the Myra Reinhard Family Foundation (JBZ), the National Institutes of Health R01NS119823 (JBZ), and the Koret Family Foundation (JBZ). ML is a Merck-sponsored fellow of the Helen Hay Whitney Foundation and a Stanford Wu Tsai Neurosciences Interdisciplinary Scholar.

---

**Supplemental Figure 1:**
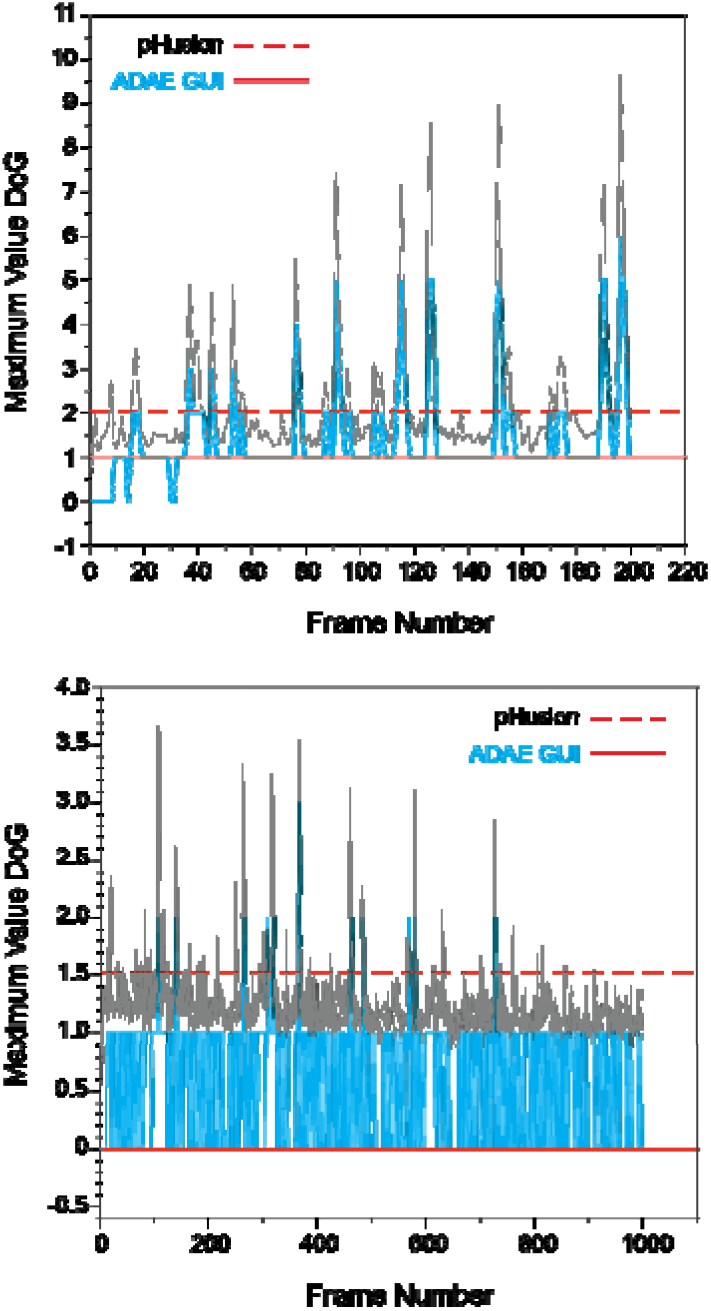
Comparison of the Difference of Gaussian (DoG) maximum value vectors calculated based on integers (ADAE GUI) or floating point numbers (pHusion) and the resulting thresholds applied. In the top panel, all of the major peaks are captured by both approaches and the resulting frequencies found are very similar: 0.062 (events/µm^2^/min), pHusion, 0.066, ADAE GUI. In the bottom panel the threshold resulting from the ADAE GUI is 0 leading to a high number of false positives and markedly different frequencies: 0.058, pHusion, 0.37, ADAE GUI. Note, to directly compare vectors between platforms the images in pHusion were converted to 8-bit images. In our standard processing pipeline images are maintained as 16-bit and thus the resulting DoG vectors are typically much greater than 1.

