## Supplementary figures and images for "pHusion: A robust and versatile toolset for automated detection and analysis of exocytosis"

### SUPPLEMENTAL FIGURE 1

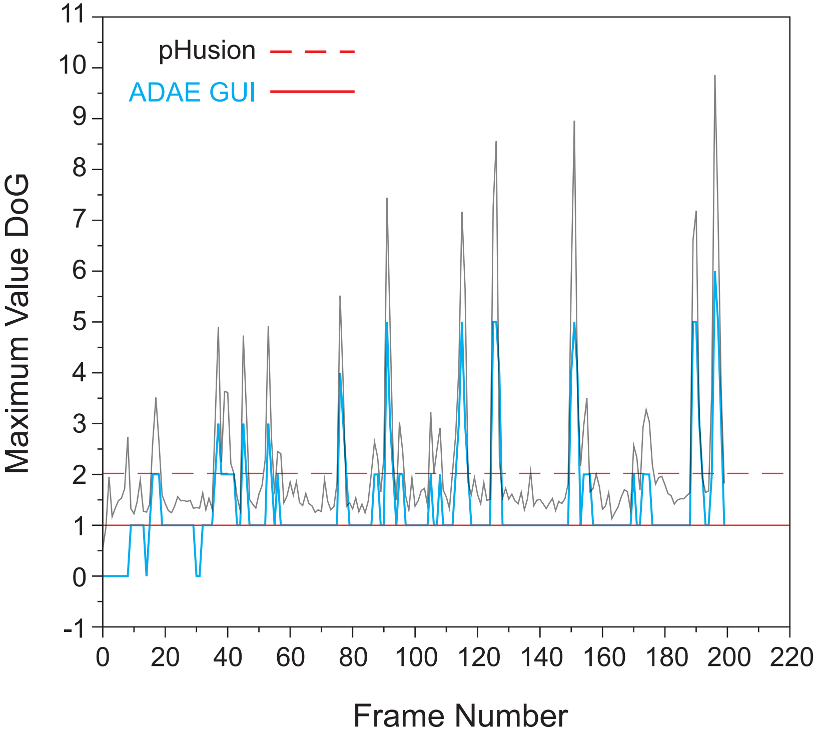

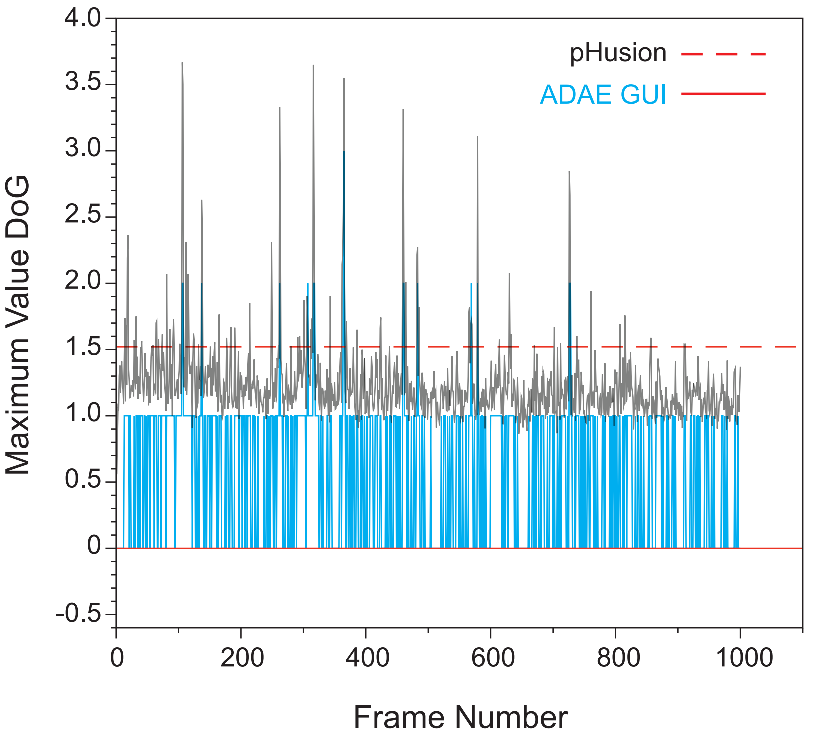
